# A reference genome without a virus: cDNA reconstruction reveals the provenance and function of the MS2 phage sequence

**DOI:** 10.64898/2026.09.18.752701

**Authors:** Eric Small, Greyson Lasley, Emily Layton, Alissa Wiwi, Devyn Del Curto, Laura D. Weinstock, Jirapat Thongchol, Junjie Zhang, Jesse Cahill

## Abstract

Reference genomes are often treated as faithful representations of experimentally validated viral genomes, yet the relationship between historically curated reference sequences and infectivity is rarely tested experimentally. Here, we developed a cDNA-based reconstruction platform for the canonical RNA phage MS2 and used it to compare the current NCBI reference genome (RefSeq) with closely related published isolate sequences. We found that isolate-derived sequences reproducibly yielded infectious phage, whereas the current MS2 RefSeq-derived construct did not, showing that the present reference does not represent a single experimentally validated infectious genome but instead reflects sequence curation across multiple studies. We then compared conventional and AI-enabled approaches to identify minimal changes that restore infectivity to MS2 RefSeq; a human experimentalist correctly prioritized corrective changes, whereas the genome language model Evo2 did not. We also observed that closely related corrected reference-derived constructs showed a ~4-log difference in phage output, and subsequent analysis indicated that this difference was associated with an apparent replicase frameshift in the lower-output background. This suggests that the low output construct class represents rare mutations from genomes that are one mutational step away from true function, rather than uniform function of the dominant construct population. A complementary cell-free assay provided a lower-background orthogonal readout of construct-level function, yielding ~1 x10^6^ PFU/mL from the high-output background within 2 hours while showing no detectable recovery from the low-output background. Together, these results establish a robust platform for RNA phage reconstruction and raise the possibility that historical reference genomes, especially for RNA viruses, may not always remain faithful to experimentally validated biological function. More broadly, these findings underscore the need to verify the infectivity of reference genomes, particularly when they were assembled non-contiguously or shaped by cumulative human curation. They also highlight the importance of clearly distinguishing historically curated reference sequences from experimentally validated infectious genomes when such data are used to train or evaluate AI/ML models.

## 1 Introduction

MS2 is one of the most historically important model viral systems in molecular biology. It is a small, positive-sense single-stranded RNA phage that infects F-pilus-bearing *Escherichia coli* and packages its ∼3.57 kb genome within a simple icosahedral capsid. That compact genome encodes only four genes—coat, maturase, replicase, and lysis—yet supports a tightly coordinated program of RNA folding, translation, replication, and assembly. Because so much biological function is compressed into such a short sequence, MS2 became an unusually accessible and modifiable system for studying how viral genomes encode infectivity, regulation, and structure [1-4].

MS2 was also the first virus to have its complete genome sequenced [5], but that milestone was reached incrementally rather than in a single experiment. Over the course of the 1970s, different portions of the genome were resolved in three steps—the coat gene first [6], then the 3′ terminal region [7], and finally the remaining portion of the genome resulting in a 3,569-nt genome assembled from overlapping RNA fragments and enzymatic sequencing studies [5]. That sequence rapidly became the gold-standard reference genome and anchored decades of work on RNA phage biology, helping establish core principles of translational control [8, 9], RNA secondary structure [10], capsid assembly [11], and RNA-dependent RNA replication [12].

Despite its central role in molecular biology, the modern MS2 reference sequence is widely used as the canonical genome sequence, but the complete reference sequence has not, to our knowledge, been directly validated as an infectious genome following de novo reconstruction. The lack of validation is concerning because of the room for error in combining legacy sequence submissions. This is compounded by the fact that RNA virus populations drift rapidly, and historically assembled sequences may not remain functionally identical to extant infectious isolates or to the material originally studied experimentally. Indeed, synthetic reconstruction has revealed functionally consequential discrepancies in other RNA virus reference sequences [13, 14]. At the same time, the historical path by which the MS2 sequence was assembled raises an important question: whether the present reference record still corresponds to a single experimentally traceable infectious genome.

This issue is especially relevant because cDNA systems for producing viable MS2 have existed for decades, yet they were generated by reverse transcription of RNA from live phage isolates [15] rather than by synthetically rebuilding and testing the reference sequence itself. Furthermore, basic platform characterization steps were left largely unaddressed by early studies. By modern standards, a fully characterized platform would be expected to define performance across cloning and expression hosts, demonstrate stability, and provide a framework for linking engineered sequence changes to infectivity.

Here we sought to address both the reference-genome question and the platform question at once. Using a modern cDNA-based MS2 reconstruction system, we asked whether the current MS2 reference sequence, when rebuilt entirely from synthetic DNA, represents an experimentally validated infectious genome, or whether its present form reflects historical sequence curation rather than a single traceable infectious isolate. At the same time, we sought to establish a more rigorously characterized MS2 reconstruction platform for testing how defined sequence changes affect infectivity. By doing so, we aimed to clarify the relationship between sequence lineage, cDNA reconstruction, and infectious phenotype in one of the most important model RNA phages. This question is especially timely because RNA-phage cDNA systems are increasingly being used not only for recovery of infectious particles, but also for genome-wide engineering and functional mapping, as illustrated by recent insertion-tolerance mapping of MS2 and PP7 [16].

## 2 Methods

### 2.1 Bacterial strains, phages, and growth conditions

MS2 and its cognate *Escherichia coli* host strain were obtained from ATCC (#15597-B1 and #15597, respectively). Cultures were propagated in LB broth or on LB agar (Teknova) at 37 °C. Overnight cultures were prepared from single colonies inoculated into 4 mL LB in culture tubes with vented caps and incubated with aeration overnight.

### 2.2 Construction and handling of MS2 cDNA plasmids

MS2 cDNA plasmids used in this study were uploaded to Genbank (PZ865118-127) and are briefly described in Table 1. Constructs were assembled from synthetic double-stranded DNA fragments obtained from Integrated DNA Technologies or Twist Bioscience using NEBuilder HiFi DNA Assembly Master Mix (NEB, #E2621) and transformed into commercially competent NEB 5-alpha *E. coli* cells (NEB, #C2987) according to the manufacturer’s instructions. Following overnight incubation at 37 °C on LB agar supplemented with 100 µg/mL carbenicillin, single colonies were inoculated into 4 mL of selective LB medium and grown overnight. Glycerol stocks were prepared from confirmed transformants and stored at 80 °C in 25% glycerol. Plasmid DNA was recovered from overnight cultures using a QIAprep Spin Miniprep Kit (Qiagen) according to the manufacturer’s recommendations.

**Table 1.** MS2 cDNA constructs used in this study. Listed are the plasmid identifiers, reader-friendly construct names, brief notes on sequence background or corrective design, and associated GenBank accessions. Constructs pJLC251–299 share the 5′ engineered expression architecture used in the second-generation platform, including upstream transcriptional control and transcript-processing features (e.g., terminator insulation, *lacO*, and hammerhead ribozyme), as described in detail in the main text. FS= frameshift caused by deletion of nucleotide t at position 2750. Figure 2A shows the approximate location of nucleotide or aa changes for each construct within the MS2 genome map, relative to the reference sequence.

| Plasmid | Alias | Notes and Features | GenBank # |
| --- | --- | --- | --- |
| pJLC194 | RefSeq | Synthesized cDNA from RefSeq NC_001417.2 | PZ865118 |
| pJLC222 | TAMU-Isolate | RT-derived cDNA from live isolate | PZ865119 |
| pJLC228 | Nearest-Isolate | Synthesized cDNA from GenBank GQ153927.1 | PZ865120 |
| pJLC251 | TAMU-Isolate | pJLC222 cDNA w/ 5’ engineering (*lacO*/HHR) | PZ865121 |
| pJLC294 | RefSeq-Min | NC_001417.2 with minimal correction | PZ865122 |
| pJLC295 | RefSeq-Min-FS | pJLC294-derived construct carrying Rep P134S + replicase FS | PZ865123 |
| pJLC296 | RefSeq-Evo2-A | NC_001417.2 with Evo2 correction attempt | PZ865124 |
| pJLC297 | RefSeq-Evo2-B | NC_001417.2 with Evo2 correction attempt 2 | PZ865125 |
| pJLC298 | RefSeq-Rescued | isogenic to pJLC295 w/ *Mat* D241H, no FS | PZ865126 |
| pJLC299 | Nearest-Isolate | pJLC228 cDNA w/ 5' engineering (*lacO*/HHR) | PZ865127 |

### 2.3 Assays for infectious MS2 produced from cDNA constructs

Plasmids were maintained as freeze-thaw-controlled DNA stocks, normalized to 10 ng/transformation in 1 to 3 µL water. Plasmid concentrations were determined using a Qubit fluorometer 2.0, and introduced into NEB 5-alpha or T7 Express (NEB #C2566) cells by heat-shock transformation. T7 express is an enhanced BL21-derived strain optimized for T7 promoter-driven protein expression. Time zero was defined as the outgrowth step, when tubes were placed in the incubator. To assess phage production from transformed cultures, samples were withdrawn at the three-hour timepoint and were clarified by brief centrifugation at high speed (∼16,000 × g) for 15-30 seconds. Immediately afterward, an aliquot of supernatant was removed without disturbing the cell pellet. Supernatants were then diluted as needed in SM buffer (10 mM Tris-HCl, 100 mM NaCl, 50 mM MgSO_4_, 0.01% gelatin, pH 7.5 and assayed for phage activity on bacterial lawns by plaque assay, as described below.

To demonstrate plasmid uptake and stable transformation efficiency for samples that did not yield viable cDNA-derived phage, aliquots of live transformed cells were removed after recovery 1 hr after outgrowth and plated on LB agar supplemented with carbenicillin and 0.2% glucose and incubated overnight at 37 °C. Serial dilutions were prepared in LB as needed to obtain countable colonies To quantify phage produced by leaky expression from the non-T7-expressing cloning strain NEB 5-alpha (used for plasmid archiving and maintenance), an aliquot of supernatant was withdrawn for plaque assays following 18–24 hr growth at 37 °C on LB supplemented with carbenicillin and 0.2% glucose.

### 2.4 Phage quantification and high titer lysate maintenance

To prepare the MS2 host for plaque assays, overnight cultures were subcultured 1:100 into fresh LB and incubated at 37°C with shaking. Aliquots were used within a defined post-subculture window (3– 7 h), during which host cells supported reproducible plating and infection assays.

Bacterial lawns for MS2 plaque assays were prepared by combining ~600 µL of same-day subcultured host cells with 6 mL of molten 0.6 % (w/v) top agar and immediately pouring onto LB agar plates. Phage titers from cDNA assays were determined by spotting 10 µL aliquots of serial dilutions, collected from the supernatant as described above onto the host lawn. Following overnight incubation at 37°C, plaques were counted from the highest dilution that yielded discrete, countable plaques.

High-titer MS2 lysates were prepared using standard full-plate plaque assays. Briefly, phage inoculum was mixed with host culture and molten top agar and overlaid onto LB agar plates. After incubation, confluent plates were harvested by adding 5 mL of SM buffer per plate and collecting the soft-agar layer with a plate scraper. Following clarification by centrifugation, the supernatant was filter sterilized and used as a positive control for cDNA-derived plaque assays.

### 2.5 Cell-free transcription/translation comparison

Cell-free assays were performed using two commercially available systems: PURExpress (New England Biolabs, E6800) and a TXTL system (myTXTL Pro Cell-Free Expression Kit #541000). Two MS2 cDNA-derived constructs were compared: pJLC295, a lower-output corrected reference-like construct that carries an expected lethal frameshift in the replicase reading frame, and pJLC298, a higher-output construct lacking that frameshift but carrying an additional maturase-gene mutation.

In PURExpress, 24 µL reactions were assembled according to the manufacturer’s general transcription/translation framework and included murine RNase inhibitor. Reactions were incubated at 37°C in an incubator and sampled at 2 hr for direct assay of phage output. In TXTL, 12 µL reactions were prepared using the system mix, template DNA, GamS, and a polymerase helper plasmid. TXTL reactions were incubated at 27°C in a thermocycler with the lid heater turned off and were likewise sampled at 2 hr for phage output assay. For both systems, we compared intact plasmid templates with PCR amplicon templates. The amplicons were designed with an added 5′ T7 promoter so they could directly serve the MS2 genome without the 5′ leak-control engineering sequences used in the plasmid constructs. DNA input was normalized to 10 ng per reaction.

### 2.6 Documentation, data handling, and statistics

All plates were photographed after incubation using either a modern smartphone camera or a Syngene G:BOX Chemi XX6 imaging system controlled with GENESYS software (Example plate data shown in Supplementary Figure 1). Colony counts, plaque counts, and associated metadata were entered into a running spreadsheet for longitudinal tracking of transformation and phage-output experiments.

GraphPad Prism 10.6.1 was used for plotting and statistical analysis of PFU and CFU measurements. Data points shown in the figures represent independent biological replicates; where indicated, technical replicates were averaged within each biological replicate before plotting. Error bars indicate standard deviation. For the construct comparisons shown in Fig. 2, where multiple construct classes were compared within the same assay framework, group differences were evaluated using Kruskal– Wallis tests with Dunn’s multiple-comparison correction. This nonparametric framework was used to provide a conservative rank-based comparison across construct classes without relying on Gaussian assumptions. For the G1 versus G2 comparisons specifically, we also examined the data using a conventional parametric framework in GraphPad Prism; both parametric and nonparametric approaches gave the same qualitative conclusion, namely that no significant pairwise G1–G2 difference was detected under the conditions tested.

For the cell-free comparisons shown in Fig. 3, where the primary questions were pairwise differences between constructs or timepoints within a given system, significance was assessed using unpaired t-tests. As a check on robustness, we also re-evaluated the Fig. 3 comparisons using the same nonparametric framework described above and obtained the same qualitative conclusions: the 2 h versus 4 h comparison within the NEB PURExpress condition remained non-significant, whereas the TXTL versus NEB PURExpress contrast remained significant.

### 2.7 Evo2-based ranking of candidate corrective mutations

The core objective was to use a genomic foundation model (Evo 2) to predict which specific mutations in a non-functional sequence were most responsible for loss of viability, or conversely, which mutations would be the most critical to correct to recover a functional phenotype.

#### 2.7.1 Sequence alignment and mutation identification

To identify genomic sequence differences, we first performed a pairwise sequence analysis using EMBOSS NEEDLE between a known functional sequence (NCBI: GQ153927.1) and the purported but non-functional reference sequence (NCBI: NC_001417.2). We identified all single-nucleotide variants (SNVs) and small insertions/deletions (indels) by performing character-by-character comparisons of the aligned sequences. We used a triplet-detection logic to group consecutive differences, allowing delineation of indels from point mutations.

#### 2.7.2 In silico generation of single-mutation variants

To isolate the functional contribution of each identified genomic difference, we generated a library of variant sequences from the functional sequence. For every identified SNV or indel, a corresponding variant sequence was constructed by introducing that specific mutation into the full functional GQ153927.1 sequence. This preserved the consistent context of the full genome sequence surrounding each mutation for the following predictions.

#### 2.7.3 Zero-shot functional prediction via Evo 2

We then followed a modified version of the zero-shot prediction notebook for BRCA1 mutations provided in the Evo 2 repository. Variant scoring was performed using the **evo2_1b_base** model scoring function. Although larger parameter models are available, the example notebook provided by the Evo 2 team used the evo2_1b_base model, and we therefore elected to begin with the same implementation. It remains possible that higher-parameter models (e.g., 7B or 40B) would achieve different performance or potentially approach the prioritization performance of a human expert.

The likelihood score of each variant and of the full functional parent sequence was calculated using the Evo 2 scoring function. The difference between each variant score and the functional parent score was then computed to generate a delta likelihood score for each candidate mutation. Under this framework, a more negative delta likelihood score relative to the functional sequence was interpreted as a less plausible or less tolerated variant. We then selected the two most negative-scoring SNVs for experimental testing of recovery potential.

## 3 Results

### 3.1 Live-isolate-derived MS2 sequences, but not the reference sequence, yield viable phage from the cDNA platform

To test whether the current MS2 reference sequence represents an infectious genotype, we adapted a cDNA-based reconstruction strategy around the F-pilus dependence of MS2 infection. Because MS2 can propagate only in F+ cells, we used an F− *E. coli* background for plasmid maintenance and phage production and then measured infectivity separately by plating clarified supernatants on an F+ host.

Earlier MS2 cDNA systems approached the problem with similar logic, except relied on temperature-controlled architecture; i.e., the pL/*cI857* system [15]. However, this configuration produced phage even without induction, with wild-type constructs yielding approximately 10^11^ PFU/mL during overnight growth even in F-cells. To reduce this pressure, we instead placed MS2 cDNA expression under T7 promoter control, such that constructs could be maintained in a T7 RNA polymerase– negative cloning strain and only expressed after transfer into an appropriate F− T7-expression host for phage production. This design separates routine plasmid maintenance from productive phage generation and, together with the p15A backbone, was intended to improve construct stability during propagation. The p15A origin was selected to provide a reasonable low-to-moderate copy balance between plasmid tractability and leak mitigation. A related strategy has previously been shown to generate infectious phage from cDNA for the *Caulobacter crescentus* RNA phage *ϕ*CB5 in *E. coli* [17]. Although MS2 and PP7 cDNA systems have recently been used for insertion-tolerance mapping, that work used replicase complementation in trans rather than the T7-driven, self-encoded MS2 recovery strategy used here, and did not directly evaluate whether the current MS2 RefSeq sequence is infectious after synthetic reconstruction [16]. To distinguish failure of phage production from simple failure of plasmid uptake, we measured both supernatant phage output at 3 hr post-outgrowth and colony-forming units from the corresponding transformed cultures at 1 hr post-outgrowth (Fig. 1A).

**Figure 1.**
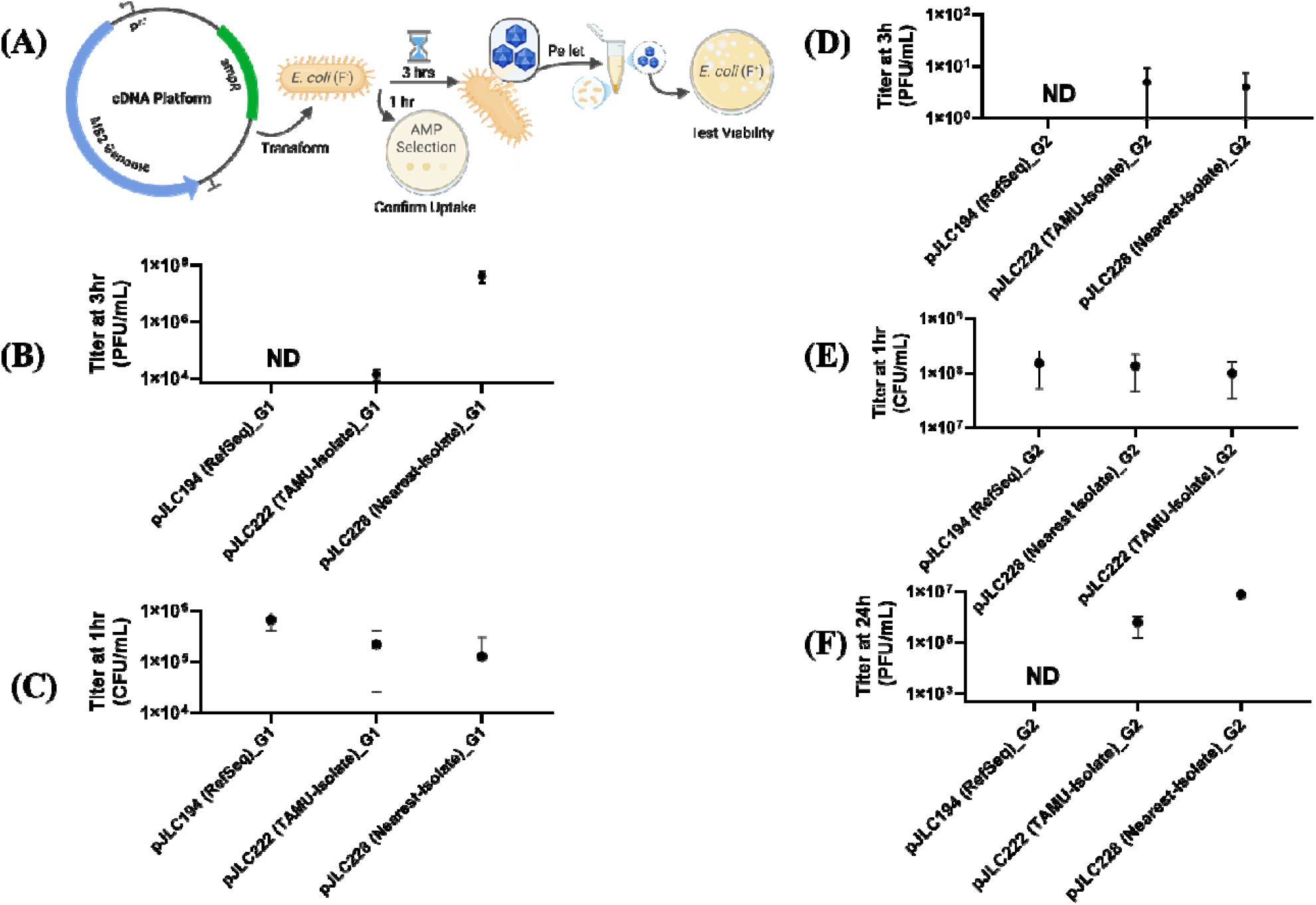
The current MS2 reference sequence does not yield viable phage in the cDNA platform, whereas isolate-derived sequences do. **(A)** Schematic of the assay used to evaluate MS2 cDNA constructs. Plasmids were transformed into F-*E. coli* host strains, and outgrowth cultures were sampled at two time points. At 1 hr post-outgrowth, aliquots of live transformed cells were removed to quantify colony-forming units (CFU) as a control for plasmid uptake and transformation efficiency. At 3 hr post-outgrowth, culture supernatants were collected and assayed for plaque-forming units (PFU) on an F+ host to quantify phage production. G1 indicates plasmid tested here after recovery following a single transformation from the original sequence-verified stock (G0). G2 indicates plasmid recovered after an additional propagation cycle from the archived strain carrying G1.**(B)** Phage production from cDNA constructs in the T7 expression system at 3 hr post-outgrowth, measured as plaque-forming units (PFU) in culture supernatants. The T7 expression system used NEB T7 Express cells, which provide T7 RNA polymerase. Induction was not required. **(C)** Transformation control for the same constructs at 1 hr post-outgrowth, measured as colony-forming units (CFU) from live transformed cells in the T7 expression system.**(D)** Performed identically to sets in panel B experiment but with G2 plasmids.**(E)** Performed identically to sets in panel C experiment but with G2 plasmids.**(F)** Free phage detected in supernatants of the non-T7 cloning strain NEB 5-alpha after overnight growth under plasmid selection, reflecting basal leakiness of the G1 constructs during plasmid maintenance in the absence of a T7 RNA polymerase expression background. Data represent 2-3 independent biological replicates, and error bars indicate standard deviation. ND= not detected.

We first tested the current MS2 reference sequence NC_001417.2 in a standard T7-driven cDNA configuration, pJLC194 (RefSeq), in which the genome was placed downstream of the canonical T7 promoter sequence TAATACGACTCACTATAGG. In this architecture, transcription initiates after the terminal TATA motif, introducing two nonviral 5′ guanosines (“GG”) into the transcript [18]. Because these promoter-proximal guanosines are commonly retained to improve T7 transcription efficiency, pJLC194 represented the most conventional implementation of the reference genome in the cDNA platform. However, pJLC194 did not yield detectable plaques in the T7 expression strain (Fig. 1B), despite showing consistent plasmid uptake at the 1 hr colony-forming unit timepoint (Fig. 1C).

We next asked whether the extra 5′ GG alone explained this failure by testing a trimmed version of the reference-derived construct that removed these promoter-derived nucleotides. That construct was also nonfunctional (data not shown), indicating that failure of the reference-derived genome could not be explained simply by the presence of the 5′ T7-associated guanosines. We therefore next tested sequences known to come from infectious MS2 isolates. The Texas A&M-derived construct, pJLC222 (TAMU-Isolate), generated previously by reverse-transcription-based cloning from a live isolate, reproducibly produced phage in the T7 expression strain (Fig. 1B). To determine whether this behavior extended beyond an internal laboratory-derived sequence, we identified the closest publicly available whole-genome MS2 isolate to the current reference sequence. By BLASTN, GQ153927.1 was the nearest whole-genome MS2 isolate in GenBank to NC_001417.2, and a construct based on this sequence, pJLC228 (Nearest-Isolate), also yielded viable phage in our system (Fig. 1B). All cDNA constructs tested are described in Table 1. Notably, the Nearest-Isolate construct produced substantially higher phage output than the TAMU-Isolate construct, a difference also detected in a related construct series below. All three constructs, including the reference-derived construct pJLC194 (RefSeq), yielded recoverable transformants at the 1 hr colony-forming unit timepoint (Fig. 1C), indicating that the failure of the RefSeq construct was not explained by lack of plasmid uptake or gross transformation inefficiency.

Next, we asked whether the same constructs retained performance after an additional cycle of archival propagation. Plasmids recovered after transformation from the original sequence-verified stock are designated G1, whereas plasmids recovered from archived strains carrying the G1 constructs are denoted G2. Compared with G1, both pJLC222 (TAMU-Isolate) and pJLC228 (Nearest-Isolate) showed markedly reduced phage output at G2 (Fig. 1D), despite continued recovery of transformed cells (Fig. 1E). In addition, the G2 plasmid preparations no longer resolved as a single clean sequence, consistent with plasmid-population heterogeneity or drift during maintenance of phage-producing constructs. In other words, continued propagation (G1-G2) imposed an additional stability burden on the system. One plausible explanation for that burden was unintended phage production during plasmid maintenance. Earlier temperature-controlled MS2 cDNA systems were known to be leaky [15], and the decline from G1 to G2 suggested that a similar problem might still be present here, even in a background designed to be free of T7 expression.

### 3.2 Platform leakiness motivates 5′ transcript-control engineering

We therefore asked whether free phage accumulated during plasmid maintenance in the non-T7-expressing cloning strain NEB 5-alpha after overnight growth on selective medium. Consistent with the T7-expression results, both pJLC222 (TAMU-Isolate) and pJLC228 (Nearest-Isolate) yielded detectable free phage in NEB 5-alpha from the G1 construct, whereas pJLC194 (RefSeq) did not (Fig. 1F). Thus, the sequence-dependent difference in phage recovery was observed both in the intended expression background and during routine plasmid maintenance.

Although the sequence-dependent difference between pJLC222 (TAMU-Isolate), pJLC228 (Nearest-Isolate) and the reference-derived construct pJLC194 was clear, the early constructs also revealed a practical limitation of the platform: the MS2 cDNA system was leaky enough that some constructs generated free phage even during plasmid maintenance in the non-T7 cloning strain NEB 5-alpha (Fig. 1F). As discussed above, unintended phage production has also been noted for earlier temperature-controlled MS2 cDNA systems [15]. This observation motivated us to test whether simple 5′ transcript-control features could suppress unintended expression during cloning while preserving productive phage recovery in the T7 expression strain. We therefore introduced commonly used design elements at the 5′ end of the cDNA cassette. The first was a lac operator (*lacO*), which provides a binding site for the LacI repressor and is widely used to reduce unwanted transcription from inducible promoters. The second was a hammerhead ribozyme (HHR), a self-cleaving RNA element placed upstream of the viral genome so that, after transcription, nonviral leader sequence is cut away to generate a cleaner 5′ RNA end. We made a third change to the platform by placing the *E. coli rrnB* T1 terminator 51 nt upstream of the T7 promoter. This was intended to insulate the MS2 cDNA cassette from run-on transcription originating elsewhere in the plasmid backbone. Plasmid numbers pJLC251-299 carry the 5’ engineering features.

### 3.3 Minimal corrective changes restore function to the reference sequence, whereas Evo2-guided corrections do not

We next tested a broader set of cDNA variants, diagrammed in Fig. 2A and described in Table 1, to better define platform behavior across related sequence backgrounds and identify the minimal corrective changes required to restore function to the reference sequence. These constructs were built in the shared 5′ engineering framework described above, allowing the effect of sequence changes to be evaluated in a common platform context. Under these conditions, the Texas A&M-derived pJLC251 (TAMU-Isolate) sequence, the publicly available isolate sequence pJLC299 (Nearest-Isolate), and the pJLC293 (RefSeq) again recapitulated the performance observed in Fig. 1: the TAMU-Isolate and Nearest-Isolate yielded viable phage, whereas the reference-derived construct did not (Fig. 2B–D).

**Figure 2.**
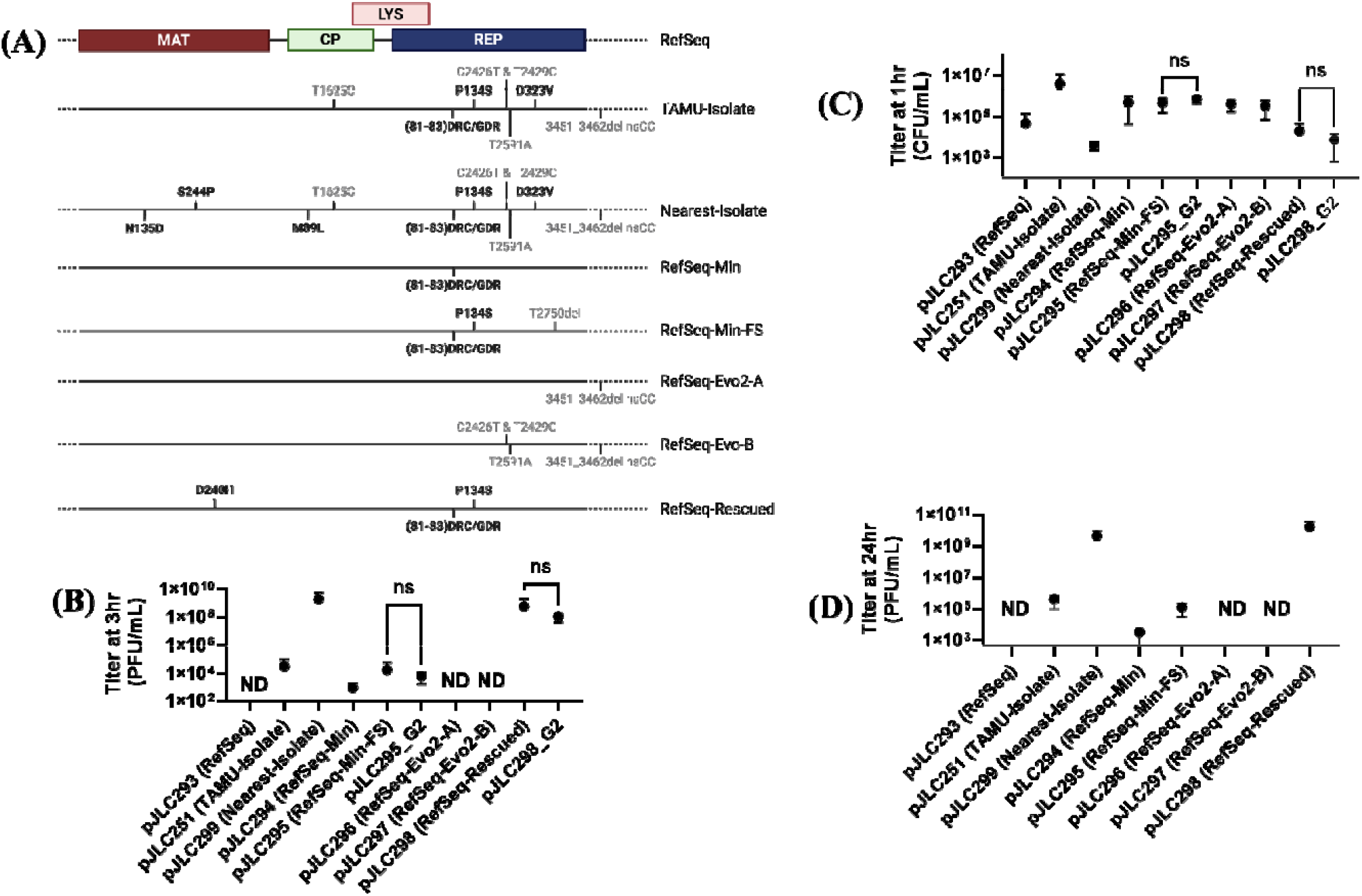
Reference-genome correction restores phage recovery in constructs carrying 5′ leak-control features. **(A)** MS2 genome maps of the constructs tested in this series, showing sequence differences relative to the current MS2 reference genome (NC_001417.2; total number of differences indicated for each construct). All constructs in this figure contained the same 5′ leak-control architecture: a lac operator (*lacO*), a hammerhead ribozyme (HHR), and an upstream *rrnB* T1 terminator positioned 51 bp before the T7 promoter. Synonymous mutations are shown as nucleotide changes in gray font, whereas non-synonymous changes are shown as amino-acid changes in black font. Unless otherwise indicated, plasmids tested in panels B–D were G1 plasmids, defined as plasmids tested after recovery from a single transformation of the original sequence-verified stock (G0). **(B)** Phage production from the indicated cDNA constructs in the T7 expression system at 3 hr post-outgrowth, measured as plaque-forming units (PFU) in culture supernatants. Where indicated, G2 denotes plasmids recovered after an additional propagation cycle from the archived strain carrying G1. **(C)** Transformation control for the same constructs at 1 hr post-outgrowth, measured as colony-forming units (CFU) from live transformed cells in the T7 expression system. Where indicated, G2 denotes plasmids recovered after an additional propagation cycle from the archived strain carrying G1. **(D)** Free phage detected in supernatants of the non-T7 cloning strain NEB 5-alpha after overnight growth under plasmid selection, reflecting basal leakiness of the constructs during plasmid maintenance in the absence of a T7 RNA polymerase expression background. All constructs shown in panels B–D were assayed using the same workflow described in Fig. 1. Points represent independent biological replicates. For G1 constructs in panels B and C, each biological replicate was measured with two technical replicates; for G2 constructs included in panels B and C, four biological replicates were measured. Error bars indicate standard deviation. For the G1–G2 comparisons, both conventional parametric and nonparametric analyses gave the same qualitative result: no significant difference was detected un er the conditions tested. **ND**, not detected.

Quantitatively, the pJLC251 (TAMU-Isolate) and pJLC299 (Nearest-Isolate) -derived constructs produced 10^4^ PFU/mL and 10^8^ PFU/mL, respectively, at 3 hr post-transformation (Fig. 2B), while corresponding transformation controls at 1 hr post-plasmid introduction showed recoverable transformants on the order of 10^4^-10^6^ CFU/mL (Fig. 2C). In general, constructs associated with higher phage output tended to yield fewer recoverable transformants, consistent with a tradeoff between productive phage generation and cellular recovery. The same sequence-dependent pattern was also observed during plasmid maintenance in the non-T7 cloning strain NEB 5-alpha (Fig. 2D). In particular, the pJLC299 (Nearest-Isolate)-derived construct and one additional high-output variant (pJLC298), described below, each reached approximately 10^9^ PFU/mL in overnight NEB 5-alpha cultures, whereas other productive constructs typically yielded on the order of 10^3^-10^5^ PFU/mL. Thus, leak-associated phage production during plasmid maintenance broadly tracked the same underlying sequence-dependent differences observed in the expression-strain assay, even though the absolute titers differed substantially between constructs.

We next asked whether the nonproductive reference-derived construct could be repaired with a minimal set of sequence changes. As one guide for this search, we leveraged Evo2, a genome language model trained on a large, diverse corpus of genomic sequence and demonstrated to support both predictive and generative biological design tasks in nucleotide sequences. We tested whether Evo2 could nominate corrective substitutions within the reference-derived genome, reasoning that if the model captured functional sequence information from its training data it would predict the substitutions most likely to recover functionality. In practice, the Evo2-guided correction predictions, tested on pJLC296-297 did not restore phage production (Fig. 2B).

In contrast, human inspection of the sequence differences between the reference-sequence construct and the viable isolate-derived constructs prioritized a small set of non-synonymous changes for testing. This strategy identified a minimal corrective set that restored viable phage output (Fig. 2A). The constructs carried on pJLC294 (RefSeq-Min) and pJLC295 (RefSeq-Min-FS) restored function to the reference sequence, with pJLC294 introducing a minimally corrective change in the replicase beginning at amino acid 80: 5′-AAT-G**<u>G</u>**T-**<u>GAT</u>**-CGC-GGT-3′, encoding N-GDR-G, changed to 5′-AAT-G**<u>A</u>**T-**<u>CGG</u>**-**<u>T</u>**GC-GGT-3′, encoding N-DRC-G. However, as defined in Table 1, pJLC295 (RefSeq-Min-FS) also carries an apparent frameshift in the replicase region caused by deletion of nucleotide T at position 2750. Although the intended Rep P134S substitution was introduced to align with the TAMU-isolate and Nearest-Isolate sequences, this frameshift was inadvertent and was expected to be lethal. This raises the possibility that the lower-output phenotype of pJLC295 does not represent a fully rescued reference background, but instead a construct one mutational step away from robust function.

This interpretation became more compelling when one pJLC295-derived clone, pJLC298 (RefSeq-Rescued), was found to lack the replicase frameshift present in pJLC295, although it also carried an additional unintended g850c mutation corresponding to a D241H substitution in the maturase protein. BLASTN searches against available *Fiersviridae* sequences indicated that the g850c change itself is not represented in closely related genomes. However, the public MS2 isolate EF204940.1 carries a different substitution at the same codon, g850a, resulting in an Asp-to-Asn change at residue 241 (D241N). This supports the idea that a negatively charged residue is not strictly required at this position.

We tested pJLC298 despite the unintended change and were surprised to find that it yielded phage production comparable to pJLC299 (Nearest-Isolate), with approximately four orders of magnitude higher PFU than the other corrected constructs in both the 3 hr T7-expression assay (Fig. 2B) and the 24 hr NEB 5-alpha leak assay (Fig. 2D). This high-output variant pJLC298 (RefSeq-Rescued) also showed a corresponding reduction in recoverable plasmid-bearing colonies at 1 hr post-transformation, consistent with a stronger burden on host recovery (Fig. 2C).

The simplest interpretation is that the constructs with lower output (pJLC294-5, pJLC251) produce phage through rare productive events, whereas the high output constructs pJLC298-9) reflect truly functional genomes (discussed later).

To test whether these constructs remained durable and sequence-stable across an additional archival propagation cycle, we examined G2 versions, with pJLC295 and pJLC298 representing a moderate-output and high-output construct, respectively. In the short-term T7-expression assay, recovered transformants and free phage production remained broadly comparable to the corresponding G1 plasmids (Fig. 2B-C), with no significant pairwise G1–G2 difference detected under the conditions tested. Both pJLC295- and pJLC298-derived plasmids could also still be successfully sequenced at G2 (Plasmidsaurus whole-plasmid nanopore sequencing). These results were striking because, despite carrying plasmids associated with phage titers approaching 10^9^ PFU/mL, the NEB 5-alpha archival strains remained recoverable and their plasmids could still be isolated and sequenced, indicating that even substantial leaky phage production did not immediately preclude plasmid maintenance. Taken together, these results show that the 5′ engineered cDNA platform is robust enough to identify functionally important defects in the MS2 reference sequence while also resolving large differences in phage output arising from relatively subtle sequence changes. Even within the highest-output class, plasmids remained recoverable, sequenceable, and assayable after an additional archival cycle, indicating that substantial leak-associated phage production does not by itself compromise short-term experimental tractability.

### 3.4 Cell-free recovery distinguishes viable from effectively nonviable MS2 cDNA genomes

One motivation for testing the MS2 cDNA system in cell-free transcription/translation was to reduce the interpretive ambiguity inherent in the in vivo assays. Plasmids are maintained within a population of living cells; therefore, rare mutations can be favored if they reduce burden or improve persistence, potentially driving drift away from the parental sequence. By contrast, a cell-free workflow removes most of that population-level maintenance and selective pressure.

To test whether the MS2 cDNA platform could be coupled to cell-free transcription/translation, we compared two systems using the same pair of constructs: a lower-output pJLC295 (RefSeq-Min-FS) and a higher-output pJLC298 (Refseq-Rescued), which lacks the replicase frameshift present in the pJLC295 background but carries a maturase point mutation. In both systems, we asked whether phage output could be recovered directly from DNA templates and whether template format affected performance.

The PURExpress system yielded substantially higher output than the TXTL system, with approximately three orders of magnitude greater phage recovery at the 2 hr timepoint for pJLC298 (RefSeq-Rescued) (Fig. 3). Additional TXTL timepoints were not pursued extensively, because our immediate goal was to identify a system capable of rapid and high-yield recovery. We also observed a strong dependence on template format. In both kits, detectable activity was recovered only when the template was supplied as a PCR amplicon carrying an added 5′ T7 promoter sequence, whereas intact plasmids did not produce comparable output under the same conditions (data not shown). This suggests that the 5′ engineering features that supported stability in vivo were not fully compatible with cell-free recovery. One possible explanation is that hammerhead ribozyme cleavage was inefficient under these conditions, thereby preventing effective generation of the intended 5’ RNA end.

**Figure 3.**
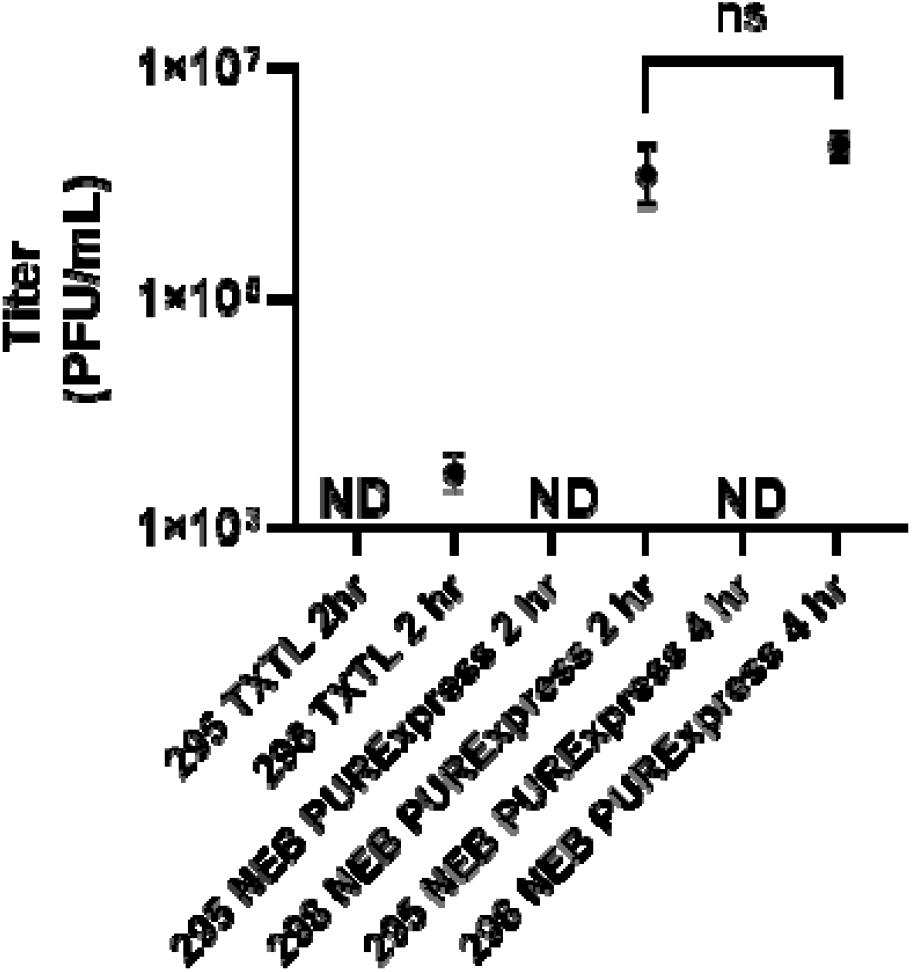
Cell-free phage recovery from two closely related MS2 cDNA amplicons differs by system and construct background. Phage output was measured as PFU/mL at the indicated timepoints for pJLC295 and pJLC298 in two cell-free systems: TXTL at 2 hr and NEB PURExpress at 2 hr and 4 hr. For each condition, PCR amplicons spanning the cDNA genome were generated from the plasmids; the amplicons included an added 5′ T7 promoter sequence, were confirmed as single bands of the expected size by gel electrophoresis, and were normalized such that 10 ng of template entered each reaction. Data is shown with the average of three independent replicates and error bars show standard deviation. Statistical comparison between the 2 hr and 4 hr timepoint for pJLC298 was performed using an unpaired t-test. ND = not detected.

Under the best-performing conditions, cell-free titers reached on the order of 1 x10^6^ PFU/mL for pJLC298 and did not increase substantially between the 2 hr and 4 hr timepoints (Fig. 3) and overnight incubation did not yield additional phage (not shown). In contrast, no detectable phage production was recovered from pJLC295 in either cell-free system, with a detection limit of

3E2 PFU/mL. Thus, even though the cell-free assays cannot exclude extremely rare productive molecules below the detection limit, the absence of detectable background provided a much clearer separation between an expected nonviable and a viable cDNA genome. In this sense, cell-free recovery may offer a useful orthogonal readout for distinguishing genuinely functional cDNA genomes from rare productive events observed in cellular systems.

## 4 Discussion

### 4.1 How prevalent is our finding in genomics?

Our central result is that the current MS2 reference sequence does not behave as an experimentally validated infectious genome in the cDNA system, whereas closely related isolate-derived sequences do. This has been observed previously for other viruses, for example, synthetic reconstruction of VSV and TMV from a reference sequence initially failed to yield virus, and subsequent experiments were able to identify sequence corrections needed to achieve rescue [13, 14]. Together, these observations suggest that reference-sequence discrepancies can have functional consequences, but they do not yet establish how frequently such discrepancies occur or how broadly they affect viral and genomic reference sequences. However, our findings combined with these identify a real and, in our view, underexplored question: how often do historically important reference genomes remain faithful to experimentally validated biology, particularly for small well-curated RNA viral genomes assembled by legacy sequencing efforts? RNA virus references may be especially vulnerable to this issue. Genome ends are biologically important and technically challenging to resolve, even with modern sequencing methods [19]. In older systems, additional uncertainty arises from how reference sequences were historically constructed. For MS2, the coat gene, terminal regions, and full genome were resolved in different studies across multiple years [5-7]. In practice, RNA phage lysates are often “refreshed” by infecting fresh host cultures with low numbers of phage and regrowing them to high titer, a process that entails many generations of infection, replication, and lysis. Given reported RNA virus mutation rates on the order of 10^-4^ substitutions per nucleotide copied, such propagation would be expected to permit measurable sequence drift over time. It is therefore unlikely that the historical sequence fragments used to assemble the canonical MS2 genome originated from the same unchanged viral population. This creates a plausible route by which the final reference could become a composite of closely related but nonidentical genomes. That possibility is not unique to MS2. Indeed, a BLASTN survey of Qbeta reference-related sequences as of August 2026 (Supplementary Figure 2) suggests that even among publicly available natural isolates, complete resolution of both 5′ and 3′ ends is often lacking. These observations do not prove that other references are non-viable, but they motivate that question as a broader investigation.

In this sense, MS2 provides a clear demonstration of a broader potential problem: gold-standard reference genomes, especially for compact RNA viruses with historically fragmented sequencing provenance, may become decoupled from experimentally validated viral genotypes. That possibility matters not only for phage biology but also for any field that treats historically curated reference sequences as faithful biological objects without direct functional confirmation.

### 4.2 cDNA platforms can be used to check reference or draft genomes

One important implication of this study is methodological: cDNA-based reconstruction platforms can be used to directly test whether reference or draft genomes correspond to viable biology. This matters even more today because the most practical route to constructing a viral cDNA system likely begins with synthetic DNA designed from publicly available genomic records, not with reverse transcription-based cloning from a live isolate [13, 14, 20]. If those records are historically curated, composite, incomplete at the ends, or otherwise functionally decoupled from viable virus, the reconstruction effort itself becomes a test of the reference rather than simply a use of it.

For MS2, earlier cDNA systems established that infectious phage could be recovered from cloned sequence, but those systems were primarily used as tools for mutational analysis rather than as fully characterized reverse-genetics platforms [15, 21]. As a result, basic questions were left largely unresolved: whether leakiness from these architectures caused toxicity in cloning strains and how genetically stable such constructs remained during maintenance in *E. coli*.

Our results suggest that cDNA reconstruction can serve not only as a tool for studying viral genomes, but also as a quality-control layer between sequence databases and experimentally validated biological function. For historically assembled reference genomes, draft viral sequences, and compact genomes, the approaches presented here offer a way to close the loop on whether a sequence corresponds to viable biology and, if not, how few changes are required to reconnect sequence to function.

### 4.3 5′ end engineering, leak mitigation, and opportunities created by leak

A major practical challenge in building an MS2 cDNA platform is controlling unintended expression in cloning strains [15]. We observed that some constructs yielded detectable free phage even during routine plasmid maintenance in NEB 5-alpha, a non-T7-expression background. Low-level, T7 RNA polymerase-independent expression has been previously reported [22, 23]. This motivated engineering of the 5′ architecture to reduce basal transcription and improve construct stability.

Specifically, we introduced a *lacO* site to recruit LacI-mediated repression, a hammerhead ribozyme (HHR) to trim nonviral 5′ transcript sequence, and an upstream *rrnB* T1 terminator to better insulate the phage cassette from plasmid-derived run-on transcription.

These additions improved the stability of the platform and preserved core sequence-dependent effects. In other words, the performance observed in Figure 1 for pJLC222 (TAMU-Isolate) and pJLC228 (Nearest-Isolate) were consistent with the 5’ engineered counterparts used in Figure 2 (pJLC251 and pJLC299, respectively). Most importantly, the performance and stability for the 5’ engineered constructs was maintained for the G2-series plasmids.

The 5′ end of the cDNA genome was a relatively conservative place to engineer the platform. MS2 replication is initiated only when the viral replicase complex recognizes the appropriate terminal sequence and RNA structural signals of authentic genomic RNA. Therefore, upstream nonviral sequence introduced for transcriptional control should have limited influence once productive replication begins, because only RNA molecules presenting the correct MS2 initiation signals at their functional genome ends would be expected to be copied efficiently. Thus, even if hammerhead ribozyme cleavage were incomplete, foreign 5′ leader sequence would be unlikely to dominate the final phenotype. In our system, the primary determinant of phage recovery was the cDNA sequence of the MS2 genome carried by the construct rather than the presence or absence of the engineered 5′ cassette.

Leakiness is therefore both a limitation and an opportunity. It complicates stable cloning and may bias recovery toward variants that suppress toxicity or reduce expression burden. In that sense, leaky maintenance is not a neutral route to generating unbiased mutant libraries; it may preferentially enrich “easy” solutions, such as single-nucleotide changes that reduce harmful expression. At the same time, this very bias could potentially be exploited in future forward-genetic experiments designed to study drift, suppression, or toxicity escape under plasmid-maintenance pressure.

### 4.4 Low-output phenotypes may reflect rare productive subpopulations

One plausible explanation for the low-output corrected constructs is that they do not represent uniformly functional sequence backgrounds but instead yield phage through rare productive subpopulations. As an order-of-magnitude estimate, rich-medium overnight cultures of *E. coli* typically contain 10□–10^1^□ CFU/mL. Assuming ~10 copies of a p15A-origin plasmid per cell [24] and a spontaneous base-substitution rate of 2 × 10□^1^□ per nucleotide per generation in wild-type *E. coli* [25], the number of plasmid copies expected to acquire a mutation at any one specified nucleotide is small, on the order of only several to tens of events under a simple approximation. The referenced mutation rate includes all possible substitutions at that site. If restoration of phage function requires one specific change, the true number of productive variants would be lower and could fall near or below the limit of detection, although local sequence context or mutational hotspots could shift this estimate upward. Nevertheless, even a small fraction of these events could generate measurable phage output. If each such cell subsequently produced 5000-10,000 phage, as reported for MS2 [26], then the above estimates of rare productive cells would be sufficient to produce 10□– 10□ PFU/mL in a 4-mL overnight culture. The estimate is therefore consistent with the low levels of free phage we observed during routine plasmid maintenance for the lower output constructs (Figure 2D). Our explanation is also consistent with recent PP7 engineering experiments in which low-efficiency recovery from an MP insertion was associated with plaque-derived genomes that had specifically eliminated the inserted sequence [16].

This interpretation also raises a second possibility: that the Texas A&M isolate sequence (pJLC251 and pJLC222), which was originally recovered by RT-PCR cloning from live phage and maintained in a higher-copy pET-style vector background (not shown), may itself have undergone subtle domestication during cloning or propagation. Even a single change that reduced burden while preserving infectivity could shift the apparent sequence background recovered from the original isolate. If so, the TAMU-derived construct may not perfectly reflect the founding viral sequence, but instead a sequence already selected for tractability in a plasmid context. Future work could address this by direct RNA sequencing of archived phage lysates, by renewed RT-PCR recovery from live phage stocks, or by plasmid-release-style assays designed to identify sequence drift that occurs during cloning and maintenance.

A related opportunity is to use cell free systems, which would allow better insulated plasmid maintenance by using promoterless cDNA constructs as archival substrates. Transcriptional control could be supplied at the assay stage as demonstrated in Figure 3, amplicons carrying an added 5′ T7 promoter were sufficient to recover phage. This could provide a cleaner way to preserve unstable genomes without requiring in vivo expression at all. Together, these possibilities emphasize that the importance of this work lies not only in resolving specific sequence backgrounds, but in the depth of characterization it provides for the MS2 cDNA platform. The system now comes with a clearer set of tools, controls, and cautions for interpreting function, instability, and construct-specific failure modes.

### 4.5 Utility of a cDNA platform

Beyond the reference-genome question, a cDNA platform opens a broader experimental space for MS2 genetics and synthetic virology. Changes that alter viability can be read out under the framework presented here, enabling reverse-genetics studies that move beyond isolated gene analysis and instead interrogate genome-wide tolerance to sequence change.

A recent genome-wide insertion-tolerance study of MS2 and PP7 further illustrates the expanding utility of RNA-phage cDNA systems [16]. Choi et al. used MuA-mediated insertion mutagenesis and cDNA-based recovery to identify insertion-tolerant regions in MS2 and PP7. That work is complementary to the present study: it demonstrates how reverse-genetics systems can support systematic RNA-phage engineering, whereas our study uses cDNA reconstruction to ask whether a historically curated reference sequence itself corresponds to an infectious genotype.

Finally, this platform raises the possibility of testing genomes that exist only in sequence space. Compact viral genomes found in metagenomic data are often disconnected from host or functional assignment. While host mapping remains a major challenge, reconstruction systems like the one described here suggest a route by which sequence-derived candidate genomes could eventually be functionally tested, helping bridge viral dark matter to host range and phenotype.

### 4.6 Prompt-to-phage: AI and small tractable genomes

Small phages may be one of the best systems for asking whether AI can move from sequence plausibility to true biological prediction. Few forms of life allow a genome to be maintained on a plasmid, manipulated systematically, and then scored in simple, low-risk assays as viable or non-viable. In that sense, phages with compact genomes provide a uniquely sharp genotype-to-phenotype testbed.

Our comparison between human-guided and Evo2-guided correction highlights how phage can serve as a tractable AI testbed. In this study, a human experimentalist, reasoning from sequence differences and functional context, identified a minimal set of corrective changes that restored the reference-derived genome, whereas Evo2-based candidate corrections did not. This does not mean genomic language models are uninformative, and several limitations should be acknowledged. In our case, we evaluated only single-difference corrective candidates, even though recovery of function may in some cases depend on a minimal combination of changes rather than any one mutation alone. We also used a zero-shot framework without task-specific fine-tuning on MS2 genotype-to-phenotype data. In addition, although larger and potentially more informative Evo2 variants are available, we began with evo2_1b_base because it is the model used in the publicly available developer workflow and represents a level of compute accessibility that is realistic for many users. By contrast, larger variants such as 7B or 40B require increasingly specialized multi-GPU or high-performance computing infrastructure and are therefore less representative of what can be routinely deployed outside well-provisioned environments. The present result is therefore informative about what can be achieved with relatively accessible tools, even if larger models could perform differently and potentially narrow the gap observed here between model-guided and human-guided correction. The cDNA platform described here therefore serves not only as a reconstruction system, but also as a means of generating the mutation-level corpus needed to train and test future models.

This result also raises a broader concern about the training data of genomic language models. Such models are built from public sequence corpora that are enriched for reference records and historically curated sequences. If those records do not always correspond to experimentally validated infectious genotypes, particularly in compact RNA viruses, then the model’s notion of sequence plausibility may be skewed toward reference convention rather than biological function. In that scenario, predictive performance could fail not because the model cannot learn sequence regularities, but because the underlying training targets are not anchored to viability.

Longer term, this raises the possibility of true “prompt-to-phage” design: AI systems that could nominate candidate phage genomes for specific purposes, such as species-specific reporting, delivery of foreign DNA or RNA into otherwise intractable organisms, or selective modulation of members of a microbiome. For such a vision to become realistic, however, two things will be required. First, models must be trained and tested against rigorous genotype-to-phenotype data which a drift tolerant platform could be used to generate. Second, the vast space of viral dark matter must be connected to host identity and biological function and recapitulated leveraging synthetic biology. In that sense, the MS2 cDNA system is not only a reconstruction tool, but also a robust proving ground for the broader question of whether sequence can be converted into predictive, purposeful phage biology.

## 5 Permission to reuse and Copyright

This is an open-access article distributed under the terms of the Creative Commons Attribution License (CC BY). The use, distribution or reproduction in other forums is permitted, provided the original author(s) and the copyright owner(s) are credited and that the original publication in this journal is cited, in accordance with accepted academic practice. No use, distribution or reproduction is permitted which does not comply with these terms.

## 7 Disclaimer

This paper describes objective technical results and analysis. Any subjective views or opinions that might be expressed in the paper do not necessarily represent the views of the U.S. Department of Energy or the United States Government.

## 8 Conflict of Interest

The authors declare that the research was conducted in the absence of any commercial or financial relationships that could be construed as a potential conflict of interest.

## 9 Author Contributions

ES: Conceptualization, Data Curation, Investigation, Methodology, Project Administration, Resources, Supervision, Validation, Writing—Original Draft, Writing—Review & Editing

GL: Data Curation, Investigation, Methodology, Resources, Validation, Visualization, Writing— Review & Editing

EML: Investigation, Methodology, Validation, Visualization, Writing—Review & Editing AW: Investigation, Resources, Writing—Review & Editing

DDC: Investigation, Resources, Writing—Review & Editing

LW: Conceptualization, Formal Analysis, Methodology, Validation, Writing—Review & Editing

JT: Methodology, Resources, Writing—Review & Editing

JZ: Methodology, Resources, Writing—Review & Editing

JC: Conceptualization, Formal Analysis, Methodology, Project Administration, Resources, Supervision, Validation, Visualization, Writing—Original Draft, Writing—Review & Editing

## 10 Funding

The author(s) declare financial support was received for the research, authorship, and/or publication of this article. This study was supported by the Laboratory Directed Research and Development program at Sandia National Laboratories. Sandia National Laboratories is a multimission laboratory managed and operated by National Technology and Engineering Solutions of Sandia, LLC, a wholly-owned subsidiary of Honeywell International Inc., for the U.S. Department of Energy’s National Nuclear Security Administration under contract DE-NA0003525. JT and JZ are supported by the Center for Phage Technology, the NIH grant R01GM141659, TAMU ADM grant. JT and JZ are co-founders of Phanetica LLC. E.M.L was supported by the Universities Research Association - Sandia Graduate Student Summer Fellowship and the Lawrence M. Blatt Biotechnology Internship Scholarship.

## Acknowledgments

We thank the Cahill Laboratory members and the Sandia National Laboratories’ Environmental Systems Biology and Molecular and Microbiology Department for their valuable input during this study. We thank Isabella Romano for reviewing a pre-submission manuscript draft, suggesting edits, and providing technical feedback. SandiaAI Chat 5.4, a version of OpenAI’s GPT-5 architecture, was used for language editing and organizational suggestions. All scientific content, interpretation, and final wording were reviewed and approved by the authors.

## Data Availability Statement

All datasets generated and analyzed during this study are included in the article and its Supplementary Material.

## Notes

### Competing Interest Statement

The authors have declared no competing interest.

### Summary of Updates

-improved resolution of figures -captured recent cDNA citation/prior work

## Works Cited

1. Thongchol, J., et al., Recent advances in structural studies of single-stranded RNA bacteriophages. Viruses, 2023. 15(10): p. 1985.

2. Dent, K.C., et al., The asymmetric structure of an icosahedral virus bound to its receptor suggests a mechanism for genome release. Structure, 2013. 21(7): p. 1225–1234.

3. Rolfsson, Ó., et al., Direct Evidence for Packaging Signal-Mediated Assembly of Bacteriophage MS2. Journal of Molecular Biology, 2016. 428(2, Part B): p. 431–448.

4. Meng, R., et al., Structural basis for the adsorption of a single-stranded RNA bacteriophage. Nature communications, 2019. 10(1): p. 3130.

5. Fiers, W., et al., Complete nucleotide sequence of bacteriophage MS2 RNA primary and secondary structure of the replicase gene. Nature, 1976. 260: p. 500–507.

6. Min Jou, W., et al., Nucleotide sequence of the gene coding for the bacteriophage MS2 coat protein. Nature, 1972. 237: p. 82–88.

7. Vandenberghe, A., W. Min Jou, and W. Fiers, 3’-Terminal nucleotide sequence (n equals 361) of bacteriophage MS2 RNA. Proceedings of the National Academy of Sciences, 1975. 1975(7): p. 2559–2562.

8. Berkhout, B., Translational control mechanisms in RNA bacteriophage MS2. 1986, Leiden, The Netherlands: Ph.D. thesis, Leiden University.

9. Valegård, K., et al., Crystal structure of an RNA bacteriophage coat protein–operator complex. Nature, 1994. 371(6498): p. 623–626.

10. Koning, R.I., et al., Asymmetric cryo-EM reconstruction of phage MS2 reveals genome structure in situ. Nature communications, 2016. 7(1): p. 12524.

11. Stockley, P.G., et al., Bacteriophage MS2 genomic RNA encodes an assembly instruction manual for its capsid. Bacteriophage, 2016. 6(1): p. e1157666.

12. Wagner, A., L.I. Weise, and H. Mutschler, In vitro characterisation of the MS2 RNA polymerase complex reveals host factors that modulate emesviral replicase activity. Communications biology, 2022. 5(1): p. 264.

13. Moles, C.M., et al., Leveraging synthetic virology for the rapid engineering of vesicular stomatitis virus (VSV). Viruses, 2024. 16(10): p. 1641.

14. Cooper, B., Proof by synthesis of Tobacco mosaic virus. Genome Biology, 2014. 15(5): p. R67.

15. Olsthoorn, R. and J. van Duin, Random removal of inserts from an RNA genome: selection against single-stranded RNA. Journal of virology, 1996. 70(2): p. 729–736.

16. Choi, S.-Y., et al., Systematic mapping of insertion-tolerant regions enables capsid engineering of an infectious RNA phage. mBio, 2026. 2026(9): p. e01639–26.

17. Wang, Y., et al., Structural mechanisms of Tad pilus assembly and its interaction with an RNA virus. Science Advances, 2024. 10(18): p. eadl4450.

18. Jiang, M., et al., Interrupting the template strand of the T7 promoter facilitates translocation of the DNA during initiation, reducing transcript slippage and the release of abortive products. Journal of Molecular Biology, 2001. 310(3): p. 509–522.

19. Ni, N. and G. Burgio, Nanopore direct RNA sequencing (DRS) of MS2 bacteriophages in E. coli throughout its life cycles reveals a complex transcriptional activity to control and maintain its growth. Virology Journal, 2026. 23(1): p. 99.

20. Del Curto, D., et al., Assessment of the phi6 lysis system using genetic complementation and heterologous expression in Escherichia coli and Pseudomonas syringae. Frontiers in Microbiology, 2025. 16: p. 1718418.

21. Klovins, J., et al., Rapid evolution of translational control mechanisms in RNA genomes. J.Molelcular Biology, 1997. 265(4): p. 372–384.

22. Lagos, R., J.E. Villanueva, and O. Monasterio, Identification and properties of the genes encoding microcin E492 and its immunity protein. Journal of bacteriology, 1999. 181(1): p. 212–217.

23. Somerville, R.L., et al., Gene expression from multicopy T7 promoter vectors proceeds at single copy rates in the absence of T7 RNA polymerase. Biochemical and biophysical research communications, 1991. 181(3): p. 1056–1062.

24. Kar, S. and A.D. Ellington, Construction of synthetic T7 RNA polymerase expression systems. Methods, 2018. 143: p. 110–120.

25. Foster, P.L., et al., Determinants of spontaneous mutation in the bacterium Escherichia coli as revealed by whole-genome sequencing. Proceedings of the National Academy of Sciences, 2015. 2015(44): p. E5990–E5999.

26. Grosjean, H. and W. Fiers, Preferential codon usage in prokaryotic genes: the optimal codonanticodon interaction energy and the selection codon usage in efficiently expressed genes. Gene, 1982. 18: p. 199–209.

