## Supplemental Content for "A reference genome without a virus: cDNA reconstruction reveals the provenance and function of the MS2 phage sequence" (from another version of this preprint: version 1, 2026-09-19)

**Supplemental Information**

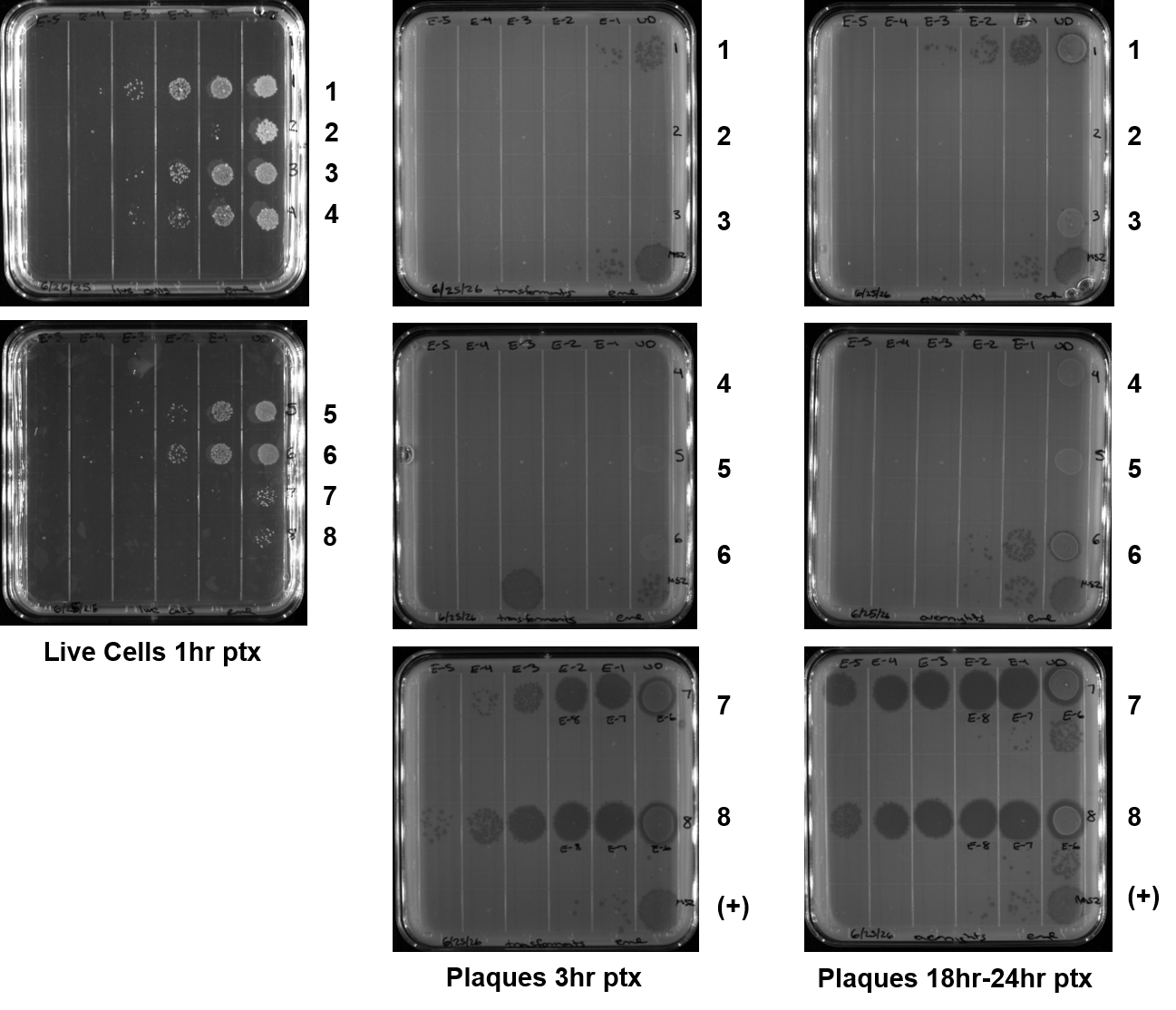

Figure S1. Representative plate images used for assay readouts in this study. Left, CFU from live transformed T7 Express cells plated on LB agar supplemented with carbenicillin at 1 hr post-transformation. Middle, PFU measured from the supernatant 3 hr post-transformation into T7 express. Right, plaques recovered after 18–24 h from NEB 5-alpha cells carrying the plasmid, showing leak-associated phage production from an MS2 wild-type construct.

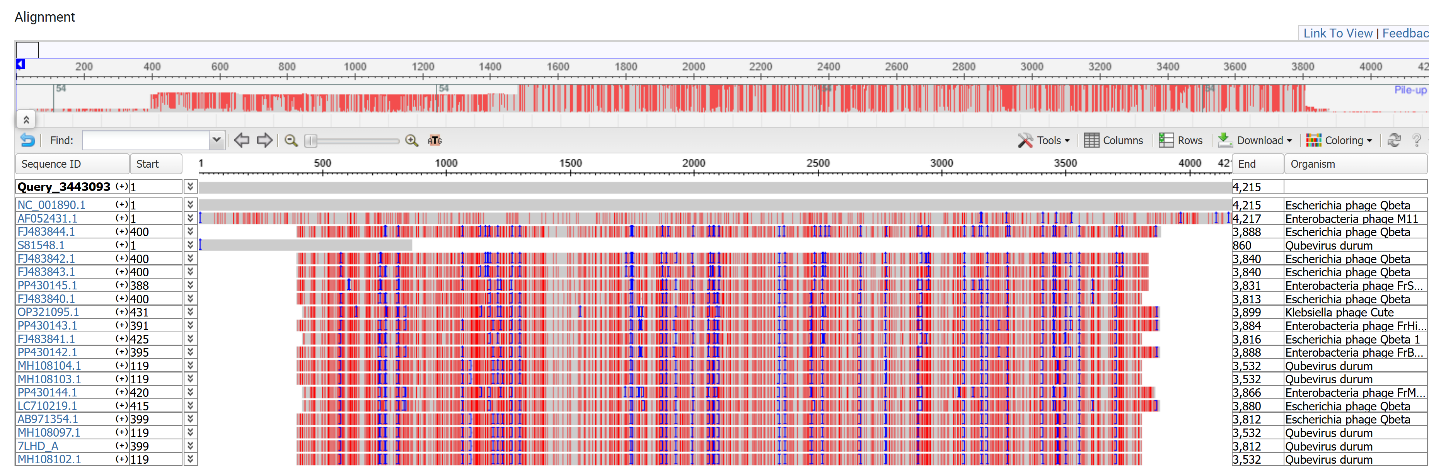

Figure S2. NCBI BLASTN multiple-sequence-alignment viewer screenshot for the bacteriophage Qß reference genome (NC_001890.1). The displayed alignment shows the nearest matching reference and isolate sequences returned by BLASTN for the Qß genome and illustrates the incomplete representation of terminal sequence information among the closest publicly available comparators.

Table S1. Primers used in this study. Listed are the primer names, their experimental purpose, and nucleotide sequences in the 5′→3′ orientation. Primers include those used for TXTL amplicon generation and for amplification of MS2 cDNA fragments or plasmid backbone components for Gibson assembly.

| Primer | Purpose | Sequence (5’-3’) |
| --- | --- | --- |
| TXTL for | Amplifies MS2 cDNA from 5’ end and adds a T7 promoter sequence | taaggataatacgactcactatagggggtgggacccctttcg |
| TXTL rev | Amplies MS2 cDNA from 3’ end | tgggtggtaactagccaagc |
| 903 331 for | Amplifies MS2 cDNA fragments for Gibson assembly obtained from IDT or Twist | gaaatctccgccccgttcgtaaATAAAACGAAAG |
| 886 331 rev | Amplifies MS2 cDNA fragments for Gibson assembly obtained from IDT or Twist | cagggttattgtctcatgagcggatacatatttg |
| 882 pJLC222 term down | Amplifies a p15A and AmpR-cassette backbone for Gibson Assembly | GAAGACTGGGCCTTTCGTTTTATttacgaacggggcggagatttc |
| 888 pJLC222 for | Amplifies a p15A and AmpR-cassette backbone for Gibson Assembly | caaatatgtatccgctcatgagacaataaccctg |

Sequences used for assembly.
Shown is an example double-stranded DNA assembly sequence based on the TAMU isolate used to generate pJLC251 (see Table S1 for primers). In the annotated sequence, the T7 promoter, *lacO* sequence, and hammerhead ribozyme are highlighted in red, blue, and purple, respectively.

GAAATCTCCGCCCCGTTCGTAAATAAAACGAAAGGCCCAGTCTTCCGACTGAGCCTTTCGTTTTATAGGAAGCGGAATATATCCCTAGGTCTAGGGCGGCGGATGAAGCGTTAAGGATAATACGACTCACTATAGGAATTGTGAGCGCTCACAATTCCGCTGATGAGGCCGAAAGGCCGAAACCCGTCGGGTGGGACCCCTTTCGGGGTCCTGCTCAACTTCCTGTCGAGCTAATGCCATTTTTAATGTCTTTAGCGAGACGCTACCATGGCTATCGCTGTAGGTAGCCGGAATTCCATTCCTAGGAGGTTTGACCTGTGCGAGCTTTTAGTACCCTTGATAGGGAGAACGAGACCTTCGTCCCCTCCGTTCGCGTTTACGCGGACGGTGAGACTGAAGATAACTCATTCTCTTTAAAATATCGTTCGAACTGGACTCCCGGTCGTTTTAACTCGACTGGGGCCAAAACGAAACAGTGGCACTACCCCTCTCCGTATTCACGGGGGGCGTTAAGTGTCACATCGATAGATCAAGGTGCCTACAAGCGAAGTGGGTCATCGTGGGGTCGCCCGTACGAGGAGAAAGCCGGTTTCGGCTTCTCCCTCGACGCACGCTCCTGCTACAGCCTCTTCCCTGTAAGCCAAAACTTGACTTACATCGAAGTGCCGCAGAACGTTGCGAACCGGGCGTCGACCGAAGTCCTGCAAAAGGTCACCCAGGGTAATTTTAACCTTGGTGTTGCTTTAGCAGAGGCCAGGTCGACAGCCTCACAACTCGCGACGCAAACCATTGCGCTCGTGAAGGCGTACACTGCCGCTCGTCGCGGTAATTGGCGCCAGGCGCTCCGCTACCTTGCCCTAAACGAAGATCGAAAGTTTCGATCAAAACACGTGGCCGGCAGGTGGTTGGAGTTGCAGTTCGGTTGGTTACCACTAATGAGTGATATCCAGGGTGCATATGAGATGCTTACGAAGGTTCACCTTCAAGAGTTTCTTCCTATGAGAGCCGTACGTCAGGTCGGTACTAACATCAAGTTAGATGGCCGTCTGTCGTATCCAGCTGCAAACTTCCAGACAACGTGCAACATATCGCGACGTATCGTGATATGGTTTTACATAAACGATGCACGTTTGGCATGGTTGTCGTCTCTAGGTATCTTGAACCCACTAGGTATAGTGTGGGAAAAGGTGCCTTTCTCATTCGTTGTCGACTGGCTCCTACCTGTAGGTAACATGCTCGAGGGCCTTACGGCCCCCGTGGGATGCTCCTACATGTCAGGAACAGTTACTGACGTAATAACGGGTGAGTCCATCATAAGCGTTGACGCTCCCTACGGGTGGACTGTGGAGAGACAGGGCACTGCTAAGGCCCAAATCTCAGCCATGCATCGAGGGGTACAATCCGTATGGCCAACAACTGGCGCGTACGTAAAGTCTCCTTTCTCGATGGTCCATACCTTAGATGCGTTAGCATTAATCAGGCAACGGCTCTCTAGATAGAGCCCTCAACCGGAGTTTGAAGCATGGCTTCTAACTTTACTCAGTTCGTTCTCGTCGACAATGGCGGAACTGGCGACGTGACTGTCGCCCCAAGCAACTTCGCTAACGGGGTCGCTGAATGGATCAGCTCTAACTCGCGTTCACAGGCTTACAAAGTAACCTGTAGCGTTCGTCAGAGCTCTGCGCAGAATCGCAAATACACCATCAAAGTCGAGGTGCCTAAAGTGGCAACCCAGACTGTTGGTGGTGTAGAGCTTCCTGTAGCCGCATGGCGTTCGTACTTAAATATGGAACTAACCATTCCAATTTTCGCCACGAATTCCGACTGCGAGCTTATTGTTAAGGCAATGCAAGGTCTCCTAAAAGATGGAAACCCGATTCCCTCAGCAATCGCAGCAAACTCCGGCATCTACTAATAGACGCCGGCCATTCAAACATGAGGATTACCCATGTCGAAGACAACAAAGAAGTTCAACTCTTTATGTATTGATCTTCCTCGCGATCTTTCTCTCGAAATTTACCAATCAATTGCTTCTGTCGCTACTGGAAGCGGTGATCCGCACAGTGACGACTTTACAGCAATTGCTTACTTAAGGGACGAATTGCTCACAAAGCATCCGACCTTAGGTTCTGGTAATGACGAGGCGACCCGTCGTACCTTAGCTATCGCTAAGCTACGGGAGGCGAATGATCGGTGCGGTCAGATAAATAGAGAAGGTTTCTTACATGACAAATCCTTGTCATGGGATCCGGATGTTTTACAAACCAGCATCCGTAGCCTTATTGGCAACCTCCTCTCTGGCTACCGATCGTCGTTGTTTGGGCAATGCACGTTCTCCAACGGTGCCTCTATGGGGCACAAGTTGCAGGATGCAGCGCCTTACAAGAAGTTCGCTGAACAAGCAACCGTTACCCCCCGCGCTCTGAGAGCGGCTCTATTGGTCCGAGACCAATGTGCGCCGTGGATCAGACACGCGGTCCGCTATAACGAGTCATATGAATTTAGGCTCGTTGTAGGGAACGGAGTGTTTACAGTTCCGAAGAATAATAAAATAGATCGGGCTGCCTGTAAGGAGCCTGATATGAATATGTACCTCCAGAAAGGGGTCGGTGCCTTTATCAGACGCCGGCTCAAATCCGTTGGTATAGACCTGAATGATCAATCGATCAACCAGCGTCTGGCTCAGCAGGGCAGCGTAGATGGTTCGCTTGCGACGATAGACTTATCGTCTGCATCCGATTCCATCTCCGATCGCCTGGTGTGGAGTTTTCTCCCACCTGAGCTATATTCATATCTCGATCGTATCCGCTCACACTACGGAATCGTAGATGGCGAGACGATACGATGGGAACTATTTTCCACAATGGGAAATGGGTTCACATTTGAGCTAGAGTCCATGATATTCTGGGCAATAGACAAAGCGACCCAAATCCATTTTGGTAACGCCGGAACCATAGGCATCTACGGGGACGATATTATATGTCCCAGTGAGATTGCACCCCGTGTGCTAGAGGCACTTGCCTACTACGGTTTTAAACCGAATCTTCGTAAAACGTTCGTGTCCGGGCTCTTTCGCGAGAGCTGCGGCGCGCACTTTTACCGTGGTGTCGATGTCAAACCGTTTTACATCAAGAAACCTGTTGACAATCTCTTCGCCCTGATGCTGATATTAAATCGGCTACGGGGTTGGGGAGTTGTCGGAGGTATGTCAGATCCACGCCTCTATAAGGTGTGGGTACGGCTCTCCTCCCAGGTGCCTTCGATGTTCTTCGGTGGGACGGACCTCGCTGCCGACTACTACGTAGTCAGCCCGCCTACGGCAGTCTCGGTATACACCAAGACTCCGTACGGGCGGCTGCTCGCGGATACCCGTACCTCGGGTTTCCGTCTTGCTCGTATCGCTCGAGAACGCAAGTTCTTCAGCGAAAAGCACGACAGTGGTCGCTACATAGCGTGGTTCCATACTGGAGGTGAAATCACCGACAGCATGAAGTCCGCCGGCGTGCGCGTTATACGCACTTCGGAGTGGCTAACGCCGGTTCCCACATTCCCTCAGGAGTGTGGGCCAGCGAGCTCTCCTCGGTAGCTGACCGAGGGACCCCCGTAAACGGGGTGGGTGTGCTCGAAAGAGCACGGGTCCGCGAAAGCGGTGGCTCCACCGAAAGGTGGGCGGGCTTCGGCCCAGGGACCTCCCCCTAAAGAGAGGACCCGGGATTCTCCCGATTTGGTAACTAGCTGCTTGGCTAGTTACCACCCAGGCTCACCTTCGGGTGGGCCTTTCTGCGGCTGAGTTTGTTTATTTTTCTAAATACATTCAAATATGTATCCGCTCATGAGACAATAACCCTG

p15A and AmpR backbone. The backbone sequence shown is the one used to construct pJLC251 (see Table S1 for associated primers)

CAAATATGTATCCGCTCATGAGACAATAACCCTGATAAATGCTTCAATAATATTGAAAAAGGAAGAGTATGAGTATTCAACATTTCCGTGTCGCCCTTATTCCCTTTTTTGCGGCATTTTGCCTTCCTGTTTTTGCTCACCCAGAAACGCTGGTGAAAGTAAAAGATGCTGAAGATCAGTTGGGTGCACGAGTGGGTTACATCGAACTGGATCTCAACAGCGGTAAGATCCTTGAGAGTTTTCGCCCCGAAGAACGTTTTCCAATGATGAGCACTTTTAAAGTTCTGCTATGTGGCGCGGTATTATCCCGTGTTGACGCCGGGCAAGAGCAACTCGGTCGCCGCATACACTATTCTCAGAATGACTTGGTTGAGTACTCACCAGTCACAGAAAAGCATCTTACGGATGGCATGACAGTAAGAGAATTATGCAGTGCTGCCATAACCATGAGTGATAACACTGCGGCCAACTTACTTCTGACAACGATCGGAGGACCGAAGGAGCTAACCGCTTTTTTGCACAACATGGGGGATCATGTAACTCGCCTTGATCGTTGGGAACCGGAGCTGAATGAAGCCATACCAAACGACGAGCGTGACACCACGATGCCTGCAGCAATGGCAACAACGTTGCGCAAACTATTAACTGGCGAACTACTTACTCTAGCTTCCCGGCAACAATTAATAGACTGGATGGAGGCGGATAAAGTTGCAGGACCACTTCTGCGCTCGGCCCTTCCGGCTGGCTGGTTTATTGCTGATAAATCTGGAGCCGGTGAGCGTGGGTCACGCGGTATCATTGCAGCACTGGGGCCAGATGGTAAGCCCTCCCGTATCGTAGTTATCTACACGACGGGGAGTCAGGCAACTATGGATGAACGAAATAGACAGATCGCTGAGATAGGTGCCTCACTGATTAAGCATTGGTAATCTAGCATAACCCCTTGGGGCCTCTAAACGGGTCTTGAGGGGTTTTTTGCAAGCACTAGTAACAACTTATATCGTATGGGGCTGACTTCAGGTGCTACATTTGAAGAGATAAATTGCACTGAAATCTAGTAATATTTTATCTGATTAATAAGATGATCTTCTTGAGATCGTTTTGGTCTGCGCGTAATCTCTTGCTCTGAAAACGAAAAAACCGCCTTGCAGGGCGGTTTTTCGAAGGTTCTCTGAGCTACCAACTCTTTGAACCGAGGTAACTGGCTTGGAGGAGCGCAGTCACCAAAACTTGTCCTTTCAGTTTAGCCTTAACCGGCGCATGACTTCAAGACTAACTCCTCTAAATCAATTACCAGTGGCTGCTGCCAGTGGTGCTTTTGCATGTCTTTCCGGGTTGGACTCAAGACGATAGTTACCGGATAAGGCGCAGCGGTCGGACTGAACGGGGGGTTCGTGCATACAGTCCAGCTTGGAGCGAACTGCCTACCCGGAACTGAGTGTCAGGCGTGGAATGAGACAAACGCGGCCATAACAGCGGAATGACACCGGTAAACCGAAAGGCAGGAACAGGAGAGCGCACGAGGGAGCCGCCAGGGGGAAACGCCTGGTATCTTTATAGTCCTGTCGGGTTTCGCCACCACTGATTTGAGCGTCAGATTTCGTGATGCTTGTCAGGGGGGCGGAGCCTATGGAAAAACGGCTTTGCCGCGGCCCTCTCACTTCCCTGTTAAGTATCTTCCTGGCATCTTCCAGGAAATCTCCGCCCCGTTCGTAAATAAAACGAAAGGCCCAGTCTTC

Data

| 1B |  |  |  |  |  |  |  |  |  |
| --- | --- | --- | --- | --- | --- | --- | --- | --- | --- |
| pJLC222 (TAMU-Isolate)_G1 | pJLC194 (RefSeq)_G1 | pJLC228 (Nearest-Isolate)_G1 |  |  |  |  |  |  |  |
| 14000 | 0 | 30000000 |  |  |  |  |  |  |  |
| 20000 | 0 | 57000000 |  |  |  |  |  |  |  |
| 8000 | 0 |  |  |  |  |  |  |  |  |
| 1C |  |  |  |  |  |  |  |  |  |
| pJLC222 (TAMU-Isolate)_G1 | pJLC194 (RefSeq)_G1 | pJLC228 (Nearest-Isolate)_G1 |  |  |  |  |  |  |  |
| 290000 | 400000 | 4000 |  |  |  |  |  |  |  |
| 370000 | 700000 | 250000 |  |  |  |  |  |  |  |
| 700 | 900000 |  |  |  |  |  |  |  |  |
| 1D |  |  |  |  |  |  |  |  |  |
| pJLC222 (TAMU-Isolate)_G2 | pJLC194 (RefSeq)_G2 | pJLC228 (Nearest-Isolate)_G2 |  |  |  |  |  |  |  |
| 8 | 0 | 2 |  |  |  |  |  |  |  |
| 2 | 0 | 2 |  |  |  |  |  |  |  |
|  |  | 8 |  |  |  |  |  |  |  |
| 1E |  |  |  |  |  |  |  |  |  |
| pJLC222 (TAMU-Isolate)_G2 | pJLC194 (RefSeq)_G2 | pJLC228 (Nearest Isolate)_G2 |  |  |  |  |  |  |  |
| 90000000 | 2.7E+08 | 2.2E+08 |  |  |  |  |  |  |  |
| 1.7E+08 | 1E+08 | 1.5E+08 |  |  |  |  |  |  |  |
| 40000000 | 90000000 | 40000000 |  |  |  |  |  |  |  |
| 1F |  |  |  |  |  |  |  |  |  |
| pJLC222 (TAMU-Isolate)_G2 | pJLC194 (RefSeq)_G2 | pJLC228 (Nearest-Isolate)_G2 |  |  |  |  |  |  |  |
| 800000 | 0 | 7000000 |  |  |  |  |  |  |  |
| 1000000 | 0 | 7000000 |  |  |  |  |  |  |  |
| 90000 | 0 | 9000000 |  |  |  |  |  |  |  |
| 2B |  |  |  |  |  |  |  |  |  |
| pJLC251 (TAMU-Isolate) | pJLC293 (RefSeq) | pJLC294 (RefSeq-Min) | pJLC295 (RefSeq-Min-FS) | pJLC296 (RefSeq-Evo2-A) | pJLC297 (RefSeq-Evo2-B) | pJLC298 (RefSeq-Rescued) | pJLC299 (Nearest-Isolate) | pJLC295_G2 | pJLC298_G2 |
| 12000 |  | 100 | 2100 |  |  | 1.8E+08 | 9E+08 | 12000 | 80000000 |
| 400 |  |  | 2500 |  |  | 41000000 | 8E+08 | 8000 | 2.1E+08 |
| 11000 |  | 200 | 300 |  |  | 37000000 | 2.5E+08 | 2200 | 80000000 |
| 6000 |  |  | 3200 |  |  | 40000000 | 3E+08 | 3000 | 60000000 |
| 8000 |  | 1400 | 2500 |  |  | 1.6E+08 | 6.7E+08 |  |  |
| 1000 |  |  | 4000 |  |  | 36000000 | 4.9E+08 |  |  |
| 200000 |  | 2000 | 120000 |  |  | 4E+09 | 1.1E+10 |  |  |
| 9000 |  |  | 2600 |  |  | 80000000 | 9E+08 |  |  |
| 2C |  |  |  |  |  |  |  |  |  |
| pJLC251 (TAMU-Isolate) | pJLC293 (RefSeq) | pJLC294 (RefSeq-Min) | pJLC295 (RefSeq-Min-FS) | pJLC296 (RefSeq-Evo2-A) | pJLC297 (RefSeq-Evo2-B) | pJLC298 (RefSeq-Rescued) | pJLC299 (Nearest-Isolate) | pJLC295_G2 | pJLC298_G2 |
| 1200000 | 20000 | 230000 | 430000 | 380000 | 250000 | 26000 | 6000 | 900000 | 3000 |
| 1200000 | 4600 | 210000 | 280000 | 240000 | 260000 | 2200 | 3500 | 500000 | 10000 |
| 2000000 | 9000 | 500000 | 310000 | 440000 | 120000 | 5100 | 3100 | 500000 | 1000 |
| 21000000 | 280000 | 600000 | 1200000 | 1000000 | 900000 | 15000 | 5000 | 1000000 | 16000 |
| 900000 | 12000 | 230000 | 400000 | 300000 | 300000 | 80000 | 2000 |  |  |
| 700000 | 5300 | 170000 | 200000 | 220000 | 140000 | 4000 | 2000 |  |  |
| 1900000 | 10000 | 1600000 | 370000 | 440000 | 170000 | 3000 | 3000 |  |  |
| 1900000 | 28000 | 580000 | 600000 | 370000 | 600000 | 20000 | 6000 |  |  |
| 2D |  |  |  |  |  |  |  |  |  |
| pJLC251 (TAMU-Isolate) | pJLC293 (RefSeq) | pJLC294 (RefSeq-Min) | pJLC295 (RefSeq-Min-FS) | pJLC296 (RefSeq-Evo2-A) | pJLC297 (RefSeq-Evo2-B) | pJLC298 (RefSeq-Rescued) | pJLC299 (Nearest-Isolate) |  |  |
| 440000 | 0 | 2800 | 270000 | 0 | 0 | 6E+10 | 1E+10 |  |  |
| 610000 | 0 | 10000 | 90000 | 0 | 0 | 2.8E+10 | 1.2E+10 |  |  |
| 340000 | 0 | 1200 | 100000 | 0 | 0 | 1.3E+10 | 5.6E+09 |  |  |
| 33000 | 0 | 5000 | 50000 | 0 | 0 | 5E+09 | 1.5E+08 |  |  |
| 1000000 | 0 | 2100 | 250000 | 0 | 0 | 1.6E+10 | 1.8E+09 |  |  |
| 600000 | 0 | 2000 | 120000 | 0 | 0 | 7E+09 | 3.2E+08 |  |  |
| 300000 | 0 | 600 |  | 0 | 0 | 20000000 | 1.9E+09 |  |  |
| 50000 | 0 | 3600 | 20000 | 0 | 0 | 6E+09 |  |  |  |
| 3 |  |  |  |  |  |  |  |  |  |
| 295 TXTL 2hr | 298 TXTL 2 hr | 295 NEB PURExpress 2 hr | 298 NEB PURExpress 2 hr | 295 NEB PURExpress 4 hr | 298 NEB PURExpress 4 hr |  |  |  |  |
| 0 | 4000 | 0 | 1500000 | 0 | 2000000 |  |  |  |  |
| 0 | 2000 | 0 | 600000 | 0 | 1700000 |  |  |  |  |
| 0 | 3330 | 0 | 1800000 | 0 | 2800000 |  |  |  |  |
